# Modeling of a negative feedback mechanism explains age-dependent genetic architecture in reproduction in domesticated *C. elegans* strains

**DOI:** 10.1101/114348

**Authors:** Edward E. Large, Raghavendra Padmanabhan, Kathie L. Watkins, Richard F. Campbell, Wen Xu, Patrick T. McGrath

## Abstract

Most biological traits and common diseases have a strong but complex genetic basis, controlled by large numbers of genetic variants with small contributions to a trait or disease risk. The effect-size of most genetic variants is not absolute, but can depend on a number of factors including the age and genetic background of an organism. In order to understand the mechanisms that cause these changes, we are studying heritable trait differences between two domesticated strains of C. elegans. We previously identified a major effect locus, caused by a mutation in a component of the NURF chromatin remodeling complex, that regulated reproductive output in an age-dependent manner. The effect-size of this locus changes from positive to negative over the course of an animal’s reproductive lifespan. Using a previously published macroscale model of egg-laying rate in C. elegans, we show how time-dependent effect-size can be explained by an unequal use of sperm combined with negative feedback between sperm and ovulation rate. We validate a number of key predictions of this model using controlled mating experiments and quantification of oogenesis and sperm use. By incorporating this model into QTL mapping, we identify and partition new QTLs into specific aspects of the egg-laying process. Finally, we show how epistasis between two genetic variants is predicted by this modeling as a consequence of unequal use of sperm. This work demonstrates how modeling of multicellular communication systems can improve our ability to predict and understand the role of genetic variation on a complex phenotype. Negative autoregulatory feedback loops, common in transcriptional regulation, could play an important role in modifying genetic architecture in other traits.

**AUTHOR SUMMARY:** Complex traits are influenced not only by the individual effects of genetic variants, but also how these variants interact with the environment, age, and each other. While complex genetic architectures seem to be ubiquitous in natural traits, little is known about the mechanisms that cause them. Here we identify an example of age-dependent genetic architecture controlling the rate and timing of reproduction in the hermaphroditic nematode *C. elegans.* Using computational modeling, we demonstrate how this age-dependent genetic architecture can arise as a consequence of two factors: hormonal feedback on oocytes mediated by major sperm protein (MSP) released by sperm stored in the spermatheca and life history differences in sperm use caused by genetic variants. Our work also suggests how age-dependent epistasis can emerge from multicellular feedback systems.

## INTRODUCTION

Most biological traits have a strong heritable, or genetic, component. There is general interest to understand the genetic basis of these traits, often by identifying the quantitative trait nucleotides (QTNs) that underlie heritable variation segregating within a population. Two decades of studies of biological traits in humans and other model organisms has made it clear that the genetic basis controlling most biological traits is incredibly complex – not only are dozens to hundreds genes involved, but non-linear effects are also at play. While the role of statistical epistasis, or the deviation from a linear model in a sampled population, is debated (1), work in model organisms have demonstrated that genetic epistasis (2) (also called biological epistasis (3) or compositional epistasis (4)) between two loci is ubiquitous, observed in fungi (5-7), plants (8-11), insects (12, 13), nematodes (14, 15), birds (16, 17), and mammals (18-21). Environment and age are also relevant covariates, influencing the effect and onset of the QTN on the phenotype. A recent survey of natural variation in gene expression in *C. elegans* identified > 900 eQTLs with time-dependent dynamics (22). While identification of QTNs remains a worthwhile and necessary goal, development of novel approaches that can integrate and simplify how large numbers of genetic variants modify phenotypic divergence represents a necessary accomplishment for the field.

GWAS and QTL mapping, two common quantitative genetics techniques, are usually unable to identify interacting genetic variants. GWAS, which can narrow down causative genetic variants to small regions, are typically underpowered to identify statistically significant epistatic interactions due to low natural allele frequencies and the large number of statistical tests that must be performed (23). QTL mapping, on the other hand, has increased power to identify interacting QTLs due to equal allele frequencies but identifies large regions in linkage disequilibrium containing thousands of potential variants (24). Due to this inherent difficulty in studying epistasis that occurs between genetic variants that segregate within a population, studies of epistasis have typically focused on laboratory induced loss-of-function mutations through mutagenesis or RNAi. It is unknown whether the mechanisms that cause epistasis in these approaches will apply in natural populations.

As a more tractable model to understand how genetic variants impact a trait, we are focusing our studies on two *C. elegans* strains, N2 and LSJ2, derived from an individual hermaphrodite isolated in 1951. Due to a reproductive system that primarily relies on self-fertilization, the population these two strains derived from was genetically identical, at which point they were separated into distinct cultures of either solid or liquid media sometime between 1957 and 1958 (**Figure 1A**) (25) and allowed to diverge. N2 was cultured for ∼15 years on agar plates while LSJ2 was cultured for ∼50 years in liquid culture. Using next-generation sequencing, we identified 94 new mutations that were fixed in the N2 lineage and 188 new mutations were fixed in the LSJ2 lineage (26). Despite this low-level of genetic diversity, a large number of phenotypic differences distinguish the two strains. A total of five QTNs have been identified in these strains, providing empirical evidence linking variation in a neuropeptide receptor activity to changes in social behavior (27), variation in sensory gene deployment with specific chemosensory responses (25, 26, 28), variation in chromatin remodeling with life history tradeoffs decisions (29), and variation in an actetyltransferase as the source of cryptic genetic variation affecting organ development (30). To date, however, these studies have focused on QTNs for large-effect QTLs and have largely ignored the role of multigenic changes.

**Figure 1.**
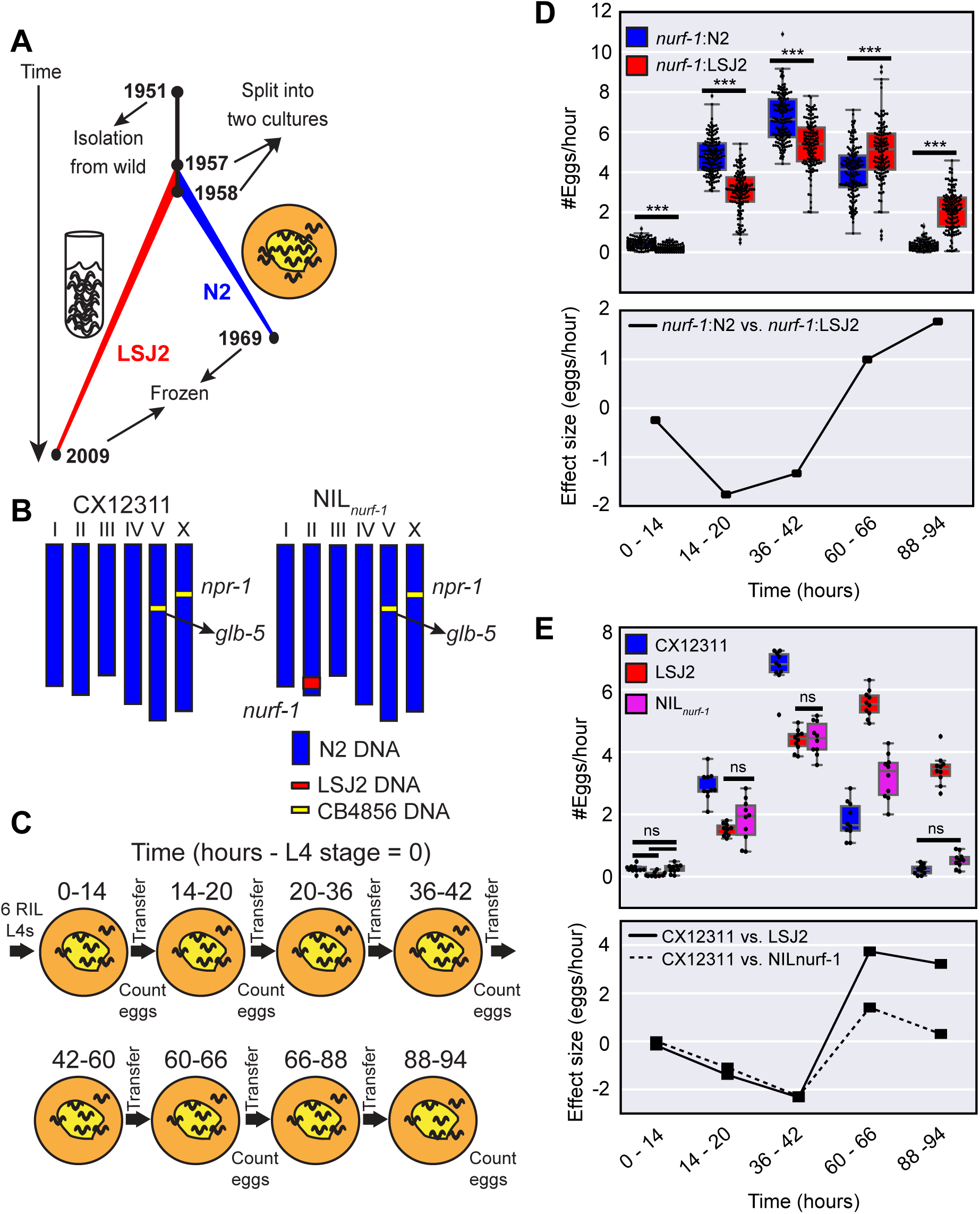
A major-effect QTL has an age-dependent effect on egg-laying **A.** History of two laboratory strains of *C. elegans* (N2 and LSJ2) following isolation of a single hermaphrodite individual from mushroom compost collected in Bristol, England in 1951. LSJ2 was grown in liquid, axenic culture whereas N2 was propagated on agar plates. **B.** Schematic of CX12311 and NIL_*nurf-1*_ strains. CB4856 is a wild strain isolated from Hawaii. N2 contains two fixed mutations in the *npr-1* and *glb-5* genes. To avoid studying their effects, we backcrossed the ancestral alleles of these genes from CB4856 into the N2 strain. *NIL_nurf-1_* contains a small region surrounding *nurf-1* backcrossed from LSJ2 into CX12311. **C**. Schematic of the experiments used to characterize the egg-laying rate at five time points. t = 0 was defined as the start of the L4 stage. **D. Top panel**. All egg-laying rate data of 94 RIL strains created between the CX12311 and LSJ2 strains. Animals were partitioned based upon their *nurf-1* genotype (blue = N2; red = LSJ2). Small difference in x-axis values for the two backgrounds are for illustration purposes only and do not indicate any difference in when the egg-laying rate was measured. Overlaid is a boxplot showing the quartiles of the data (the box) with the whiskers extended to show the rest of the distribution except for points that are determined to be outliers. All scatterplots and boxplots in the subsequent figures were calculated in the same way. For all figures, ns p >0.05, * p < 0.05, ** p < 0.01, *** p < 0.001 by Mann-Whitney U test with Bonferroni correction. **Bottom panel.** Effect size of the *nurf-1* locus measured from the RIL strains. **E. Top panel.** Egg-laying rate of CX12311, LSJ2, and NIL_*nurf-1*_ strain measured at five time points. For just this figure, only non-significant differences are shown. All other comparisons are significant at p < 0.05. **Bottom panel**. Effect size of the *nurf-1* locus measured from the NIL and parental strains.

To address this deficiency, we studied the genetic basis of reproductive differences between the N2 and LSJ2 strains at five different time points spanning their reproductive lifespan. Our goal for this study was to identify examples of complex genetic architecture and understand their molecular and cellular causes.

## RESULTS

### A major effect QTL surrounding *nurf-1* has an age-dependent effect size

We previously performed QTL mapping on reproductive rate using 94 recombinant inbred strains (RILs) generated between LSJ2 and CX12311 (29). CX12311 is a strain that derives the majority (>99%) of its DNA from N2 with the exception of a small amount of DNA backcrossed from the CB4856 wild strain that surrounds the *npr-1* and *glb-5* genes (**Figure 1B**) (26). Novel mutations in these two genes became fixed in the N2 lineage and result in pleiotropic effects on a large number of phenotypes. Use of the CX12311 strain allows us to avoid studying their effects. To study the role age plays on reproduction, we chose five time points that spanned the reproductive lifespan of the CX12311 animals. Egg-laying rate was quantified by counting the number of eggs laid by six animals for six hours on agar plates seeded with *E. coli* bacteria (**Figure 1C**). We previously identified a major effect QTL centered over the *nurf-1* gene that accounted for ∼50% of the observed phenotypic variation (26). To study how the animal’s age affected the effect-size of this locus, we first segregated the 94 RIL strains based upon their genotype at *nurf-1* (**Figure 1D – top panel**). At the first three time points, RIL strains with the N2 genotype laid more eggs than RIL strains with the LSJ2 genotype, however, at the fourth and fifth time point, this relationship flipped - animals with the LSJ2 allele of *nurf-1* laid more eggs than animals with the N2 allele. To visualize this effect more clearly, we also plotted the effect-size of the *nurf-1* locus at all five time points (**Figure 1D – bottom panel**) by subtracting the egg-laying rate of the strains with each genotype. The effect-size of the LSJ2 allele of *nurf-1* was negative for the first three time points and positive for the last two time points. This is a clear example of age-dependence, i.e. the prediction of the effect of the *nurf-1* locus on the egglaying rate requires knowledge of both the *nurf-1* genotype as well as the current animal’s age.

To verify these observations, we next assayed a near isogenic line (NIL) constructed by backcrossing the region surrounding *nurf-1* from LSJ2 into the CX12311 strain (**Figure 1B**) along with the CX12311 and LSJ2 parental strains. The CX12311 strain (containing the N2 allele of the *nurf-1* locus) laid more eggs than the NIL_*nurf-1*_ strain for the first three time points but fewer eggs at the fourth and fifth time points, again resulting in a time-dependent effect size that flips from positive to negative (**Figure 1E – top and bottom panel**). The effect size of *nurf-1* calculated using both the RIL and NIL strains and showed good qualitative agreement (**Figure 1D and 1E – bottom panel**). Due to additional segregating variants in the RIL strains, we do not expect these two calculations to be identical. In both experimental designs, the crossing of the two lines was due to a decrease in the egg-laying rate in the strain containing N2 *nurf-1* as opposed to an increase in egg-laying of the LSJ2 *nurf-1* strains. The LSJ2 strain was statistically indistinguishable at the first three time points from the NIL strain. However, at the last two time points, LSJ2 laid additional eggs resulting in a larger effect-size at these two points. These results demonstrate that the *nurf-1* locus can have both positive and negative effects on egg-laying in a manner correlated with the animal’s age.

### Difference in early egg-laying rate caused by differences in germline stem cell production, oocyte maturation and/or rate of fertilization

The rate that eggs are laid on an agar plate is dependent on a large number of factors: size and rate of mitosis of a germline progenitor pool, speed of meiosis/differentiation of these cells, maturation and growth of oocytes, ovulation and fertilization of the primary oocyte to produce an egg, and finally the rate of active expulsion of an egg through a controlled motor program that results in the opening of the vulva (**Figure 2A**). To characterize which factor might be affected by *nurf-1,* we first characterized the number of fertilized eggs in each strain using DIC microscopy. If the rate fertilized eggs were laid was affected in LSJ2 and NIL_*nurf-1*_ strains but the rate of production of fertilized eggs remained the same, we would expect LSJ2 and NIL_*nurf-1*_ to contain more eggs in their uterus than the CX12311 strain. We measured the number of fertilized eggs in each of these three strains at 24 and 48 hours after the L4 stage (**Figure 2B**). In contrast to this prediction, we found that both LSJ2 and NIL_*nurf-1*_ had significantly less fertilized eggs than the CX12311 strain. We conclude that production of fertilized eggs must be affected in the LSJ2 and NIL_*nurf-1*_ strain. The lower number of unlaid fertilized eggs in these strains is potentially a consequence of the reduced rate of production of fertilized eggs.

**Figure 2.**
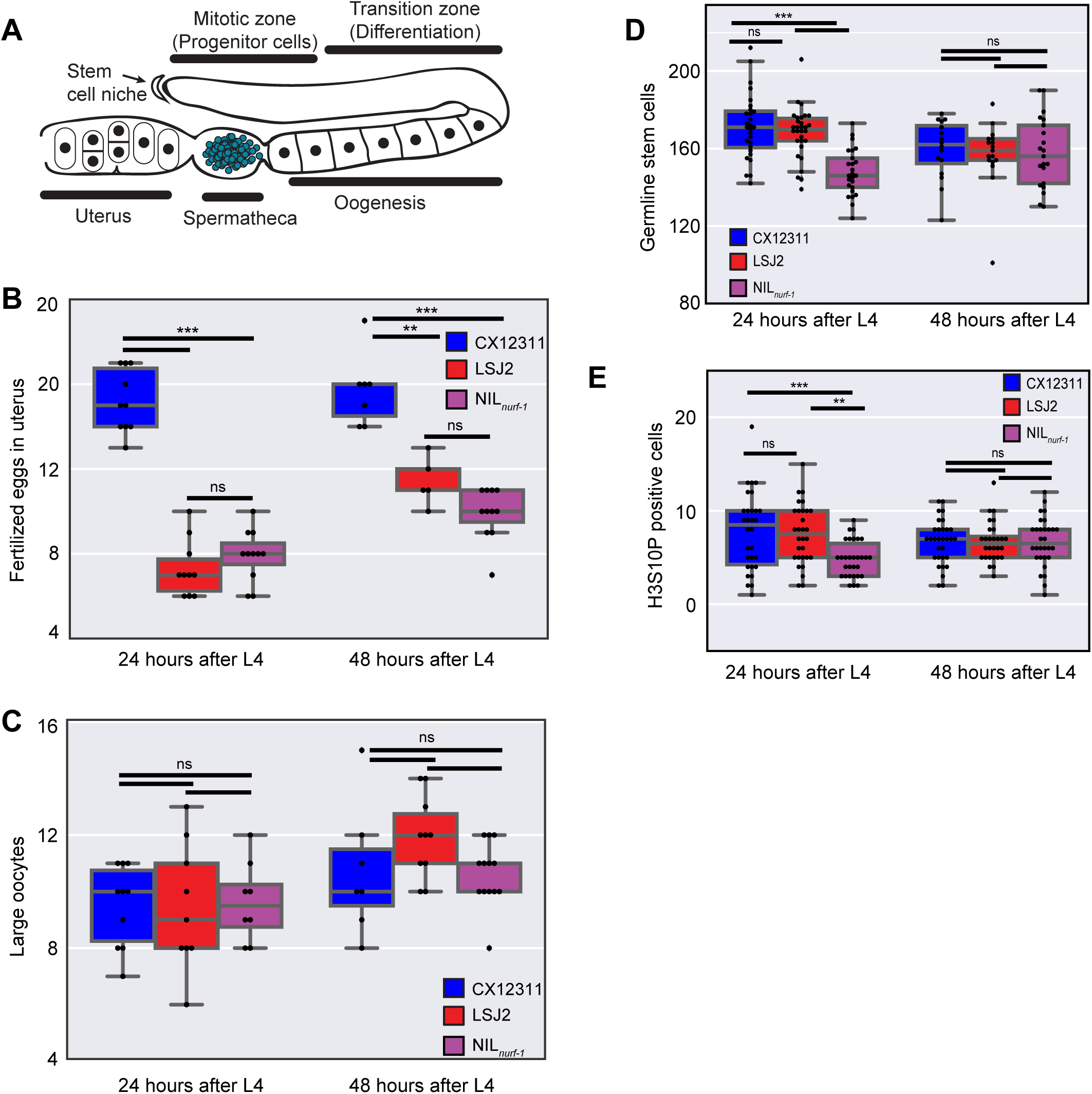
Analysis of components of egg-laying. **A.** Schematic of the *C. elegans* gonad. Germline Stem Cells (not shown due to their large number) self-renew in the mitotic zone. As they migrate away from the stem cell niche, they undergo meiosis and differentiate into mature oocytes. Ovulation forces the primary oocyte into the spermatheca, which stores previously produced self-sperm, where it is fertilized and develops an eggshell. Fertilized eggs develop in the uterus until they are laid through the vulva. Only one of two gonads is shown. **B**. Number of fertilized eggs in the uterus as determined by DIC microscopy. **C.** Number of large oocytes as determined by DAPI staining and fluorescent microscopy. **D.** Number of germline progenitor cells. **E.** Number of cells undergoing mitosis in the mitotic zone, as determined by immunofluorescence to a post-translational modification (H3S10P) in Histone 3 that is correlated with chromatin condensation in mitosis.

We next measured the number of large oocytes undergoing oogenesis in these three strains (region marked oogenesis in **Figure 2A**). We hoped to distinguish between two possible mechanisms modifying the rate of production of fertilized eggs: 1) the rate of production of mature oocytes was decreased in LSJ2/NIL_*nurf-1*_ strains or 2) the rate of fertilization of a mature oocyte was decreased in the LSJ2/NIL_*nurf-1*_ strains. We reasoned that in the former case, the number of large oocytes undergoing oogenesis would be lower in LSJ2 and NIL_*nurf-1*_ due to the decrease in production. In the latter case, if oocyte production was unaffected but the rate they were fertilized was lowered, we reasoned that the number of mature, large oocytes would increase over time. However, when we measured the number of large oocytes in each strain at 24 and 48 hours, we found no statistically significant difference between any of the three strains (**Figure 2C**). We believe this likely indicates the presence of a homeostatic mechanism that keeps the number of oocytes undergoing maturation constant, despite a difference in how often they are created or fertilized.

Finally, we measured the rate that progenitor germline stem produce new cells through mitosis, as these cells provide the source of meiotic cells that later differentiate into oocytes (marked as mitotic zone in **Figure 2A**). There is not a one-to-one relationship between the rate of mitosis and oocyte production due to cannibalization of a subset of these cells, but it is thought that there is a relationship between the size of the progenitor pool and subsequent rate of reproduction (31). Progenitor cells can be distinguished from cells in the transition zone based upon nuclear morphology. We observed a significant difference between CX12311 and NIL_*nurf-1*_ at the 24 hour time point in the number of progenitor cells, suggesting that the difference in egg-laying rate could be caused by this difference (**Figure 2D**). However, the NIL_*nurf-1*_ strain was also significantly different from LSJ2 at this time point. This difference disappeared at the 48 hour time point, when each strain had a very similar average number of progenitor cells. We also measured the number of cells undergoing mitosis in the progenitor cells, in case there was also a difference in the ratio of cells dividing in the progenitor zone. We used an antibody to Ser10 phosphorylation in histone H3, which is correlated with chromosome condensation in mitosis (**Figure 2E**) (32, 33). We observed very similar results to the germline progenitor pool suggesting the ratio of progenitor cells undergoing mitosis was the same in all three strains. The NI*L_nurf-1_* was different from both parental strains at the 24 hour time point, and all three strains were indistinguishable from each other at the 48 hours. We conclude from these observations that *nurf-1* likely has a modulatory role on the rate of early proliferation of the germline and LSJ2 contains additional genetic variations that can suppress this effect. While this difference in proliferation could account for some of the differences in egg-laying rate, we don’t believe it can explain the entire change as this would indicate different mechanisms are at play between LSJ2 and NIL_*nurf-1*_.

### A macroscale model of egg-laying can predict the age-dependent effect of the *nurf-1* locus

While age-dependent QTLs have been identified through genetic mapping approaches, few mechanisms have been demonstrated or even proposed to understand how animal’s age could influence the effect of a genetic variant. To explore possible mechanisms that could cause age-dependence for *nurf-1,* we leveraged the existing literature on genes controlling egg-laying in *C. elegans.* One recently described mechanism describes how sperm regulates the rate of egg-laying rate. *C. elegans* hermaphrodites produce a limited number of sperm (∼300) that are stored in an organ known as the spermatheca before oogenesis begins (34). Sperm release a hormone called major sperm protein (MSP) that stimulates maturation and ovulation of the primary oocyte through ephrin-receptor signaling (**Figure 3A**) on the oocyte and the gonadal sheath cell (35, 36). The effect of MSP is dose-dependent, so that as the number of sperm stored in the spermatheca decreases (through fertilization), ovulation and oocyte maturation also will decrease (35, 36). This suggests a possible mechanism tying an animal’s age to its egg-laying rate. As an animal ages, each egg it creates through fertilization reduces the number of sperm by one. Animals with N2 *nurf-1* lay more eggs during the first three time points compared to LSJ2 *nurf-1* animals and will consequently have fewer sperm at the fourth and fifth time point. This reduction in sperm will lower sperm hormone present at the primary oocyte, which counterbalances the positive effect of the N2 *nurf-1* allele on egg-laying during the transition from between the third and fourth timepoint.

**Figure 3.**
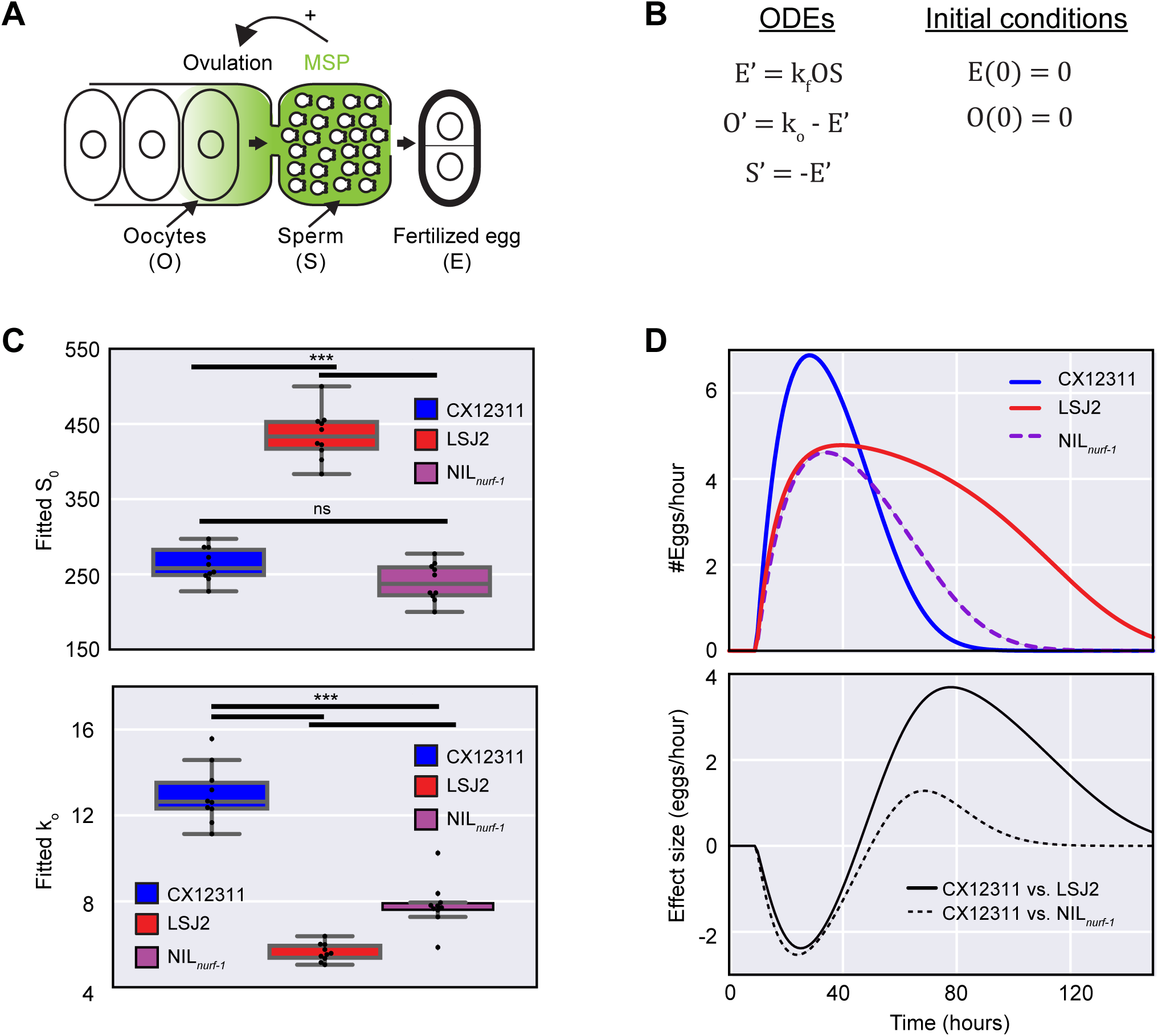
Macroscopic modeling of the effect of the *nurf-1* locus on egg-laying. **A.** Illustration of negative feedback in the egg-laying process in *C. elegans*. A limited number of sperm (200 - 350) are initially created and stored in the spermatheca before the gonad switches to exclusively produce oocytes. Sperm release a hormone called MSP, which induces oocytes to ovulate and enter the spermatheca where they are fertilized and exit into the uterus to be laid. **B.** Published macroscopic model of the egg-laying process. Ordinary differential equations (ODEs) describing the relationship between fertilized eggs (E), sperm (S), and oocytes (O) while initial conditions are given on the left. A prime (‘) indicates a time derivative. **C.** Fit of the model from figure 2B to the data plotted in figure 1E using a common value of the k_f_ parameter for all samples (0.00026). Top value shows best fit values for S_0_. Bottom panel shows best fit values for k_o_. **D**. Predicted egg-laying rate and effect-size calculated for CX12311, LSJ2, and NIL_*nurf-1*_ strains using the average value of the parameters plotted in figure 2C. This model is able to account for the rise and fall of egg-laying rate and the change in effect-size over time.

We tested this hypothesis more formally using a previously published macroscale model (37), which stipulates the egg-laying rate is proportional to the product of the number of oocytes with the number of sperm (**Figure 3B**). This model is defined by four time-independent parameters: k_o_, which specifies how rapidly oocytes are created, k_c_, which specifies a carrying capacity of the gonad, k_f_, which defines the fertilization rate, and S_0_, which specifies the number of self-sperm created by the animal. In this report, we excluded the k_c_ parameter, as we found that the carrying capacity of all three strains was similar (**Figure 2C**), and our attempts to fit this parameter resulted in negative, non-physiological values. Also, due to the nature of the equations, the k_f_ and k_o_ parameters end up having a very similar effect on egg-laying rate. To prevent the noise caused by their tight correlation, we fit a single k_f_ value to be used for all of the strains. This k_o_ value does not distinguish between the number of molecular and cellular processes described above that influence how fast mature oocytes are produced. We fit individual k_o_ and S_0_ parameters to each of the replicates in Figure 1E (**Figure 3C**). This model could recapitulate the rise and fall of the egg-laying rate - the rate rose as oocytes were generated and fell as number of sperm decreased (**Figure 3D – top panel**). A significant increase in S_0_ was observed in the LSJ2 strain compared to the CX12311 and NIL_*nurf-1*_ strain (**Figure 3C – top panel**). This is in good qualitative agreement with our previous demonstration that LSJ2 animals lay more eggs over the course of their lifetime than CX12311 (29). However, the quantitative agreement was rather poor for the LSJ2 strain. While the predicted S_0_ value of CX12311 was in good agreement with our previously measured fecundity (263 vs. 256), the predicted value of S_0_ was significantly higher than the measured fecundity (434 vs. 319). This is most likely due to the inability of the model to fully account for the genetic changes in the LSJ2 strain. The average residuals for the LSJ2 strain were significantly higher than CX12311 (2.0 vs. 0.72). The fitted k_o_ parameters matched our expectations - CX12311 had a higher k_o_ value than either the LSJ2 or NIL_*nurf-1*_ strain (**Figure 3C - bottom panel**). The effect-size calculated from the modeling experiments (**Figure 3D – bottom panel**) also qualitatively matched the effect size we experimentally measured (**Figure 1E – bottom panel**).

This analysis showed that a constant change to oocyte generation rate could have a time-dependent effect on egg-laying resulting in sign-switching at later time points. Our model explicitly predicts that the reduction in CX12311 animals in egg-laying rate at later time points is not due to changes in reproductive capacity but rather decreases in sperm number. To test this prediction, we first measured the number of fertilized eggs and large oocytes in CX12311 animals at 66 hours (**Figure 4A**). In line with our expectations, we observed a reduction in the number of fertilized eggs (∼10) contained in the CX12311 animals compared to earlier time points (**Figure 2B**). This indicates that the decrease in egg-laying rate cannot be explained by reduction of in the rate of laying fertilized eggs in the uterus. We also observed a large number of large oocytes (∼22) in CX12311 animals, an increase over the previous two time points (**Figure 4A and 2C**). This suggests that the change in egg-laying rate is not due to changes in oocyte maturation rate, as mature oocytes are available for fertilization. These observations are consistent with our hypothesis that decreased sperm and MSP at later timepoints are responsible for the decreased rates of egg-laying in CX12311.

**Figure 4.**
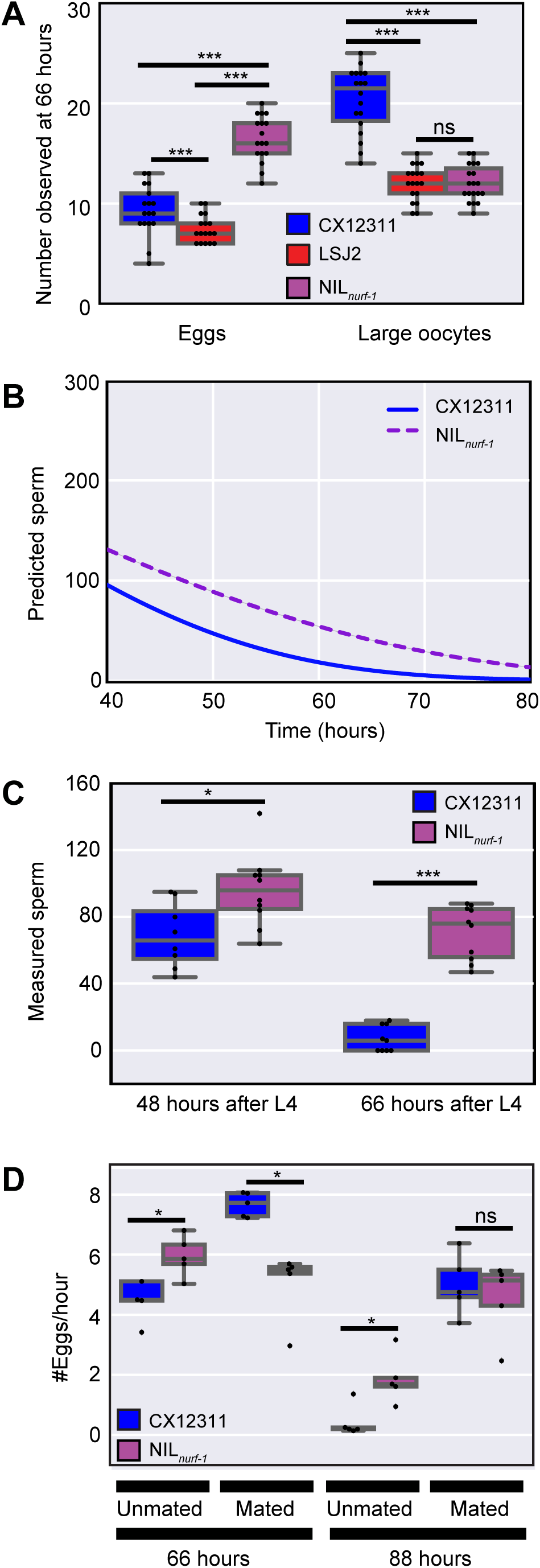
Reduction in egg-laying rate at later time points caused by use of self-sperm. **A.** Number of fertilized eggs and large oocytes at the 66 hour time point as determined by DIC microscopy and DAPI straining. This analysis indicates that CX12311, which shows a drop in egg-laying rate at this time point, is not retaining fertilized eggs in the uterus nor displaying a loss in oocytes. **B**. Modeling (identical to Figure 3) of the remaining number of self-sperm predicts that CX12311 will have fewer self-sperm than NIL_*nurf-1*_ at later points in life. **C**. Measurement of the remaining number of sperm in CX12311 and NIL_*nurf-1*_ as determined by DAPI staining and fluorescence microscopy. These results are consistent with the predictions from panel B. **D.** Egg-laying rate of CX12311 and *NIL_nurf-1_* hermaphrodites mated or unmated to CX12311 males. Mating increases the number of sperm stored in the spermatheca. These results indicate that the sign-switching observed in unmated animals (48 hours) can be reversed by the addition of sperm. The decrease in egg-laying rate in both strains at later timepoints (88 hours) can also be reversed by addition of sperm.

Our modeling predicts that the CX12311 strain will have fewer sperm later on in life due to their increase rate of fertilization at earlier time points (**Figure 4B**). We counted the number of sperm cells present in the spermatheca in CX12311 and NIL_*nurf-1*_ animals using DAPI staining combined with fluorescence microscopy. At both 48 and 66 hours, an increased number of sperm were present in the NIL_*nurf-1*_ animals compared to CX12311 in a manner consistent with the modeling.

As a final method to test this model, we took advantage of *C. elegans’* androdioecious mating system to increase the sperm available to the hermaphrodites at later time points by mating CX12311 and NIL_*nurf-1*_ hermaphroties to CX12311 males, which result in the transfer of ~1000 sperm to the hermaphrotite. After mating young adult animals, we separated the hermaphrodites from males and measured the egg-laying rate at two time points late in life. Consistent with our predictions, increasing sperm number prevented the sign-switching from occurring at both of these times in unmated animals (**Figure 4D**). This indicates that decreases in sperm number are the primary reason for decreases in egg-laying rate at late time points.

### Identification of modifier QTLs that also effect the egg-laying process

We next investigated whether additional QTLs affected egg-laying rate differences between LSJ2 and CX12311. The presence of a major effect QTL (such as the *nurf-1* locus) can mask the effects of smaller QTLs so we performed additional scans using the genotype of *nurf-1* as an additive or interacting covariate (**Figure 5A**) (38). We identified five QTLs that were significant at the genome-wide level: one QTL on the center of chromosome I, one QTL on the center of II, one large QTL on the center and right arm of chromosome IV, one QTL on the right arm of V, and one QTL in the center of × (**Figure S1 – S5**). The identification of the QTL on chromosome I was expected, as it contains a previously described missense mutation in *nath-10* that was shown to affect egg-laying (30). However, the other four QTLs do not contain any genetic variants that are known to affect egg-laying.

**Figure 5.**
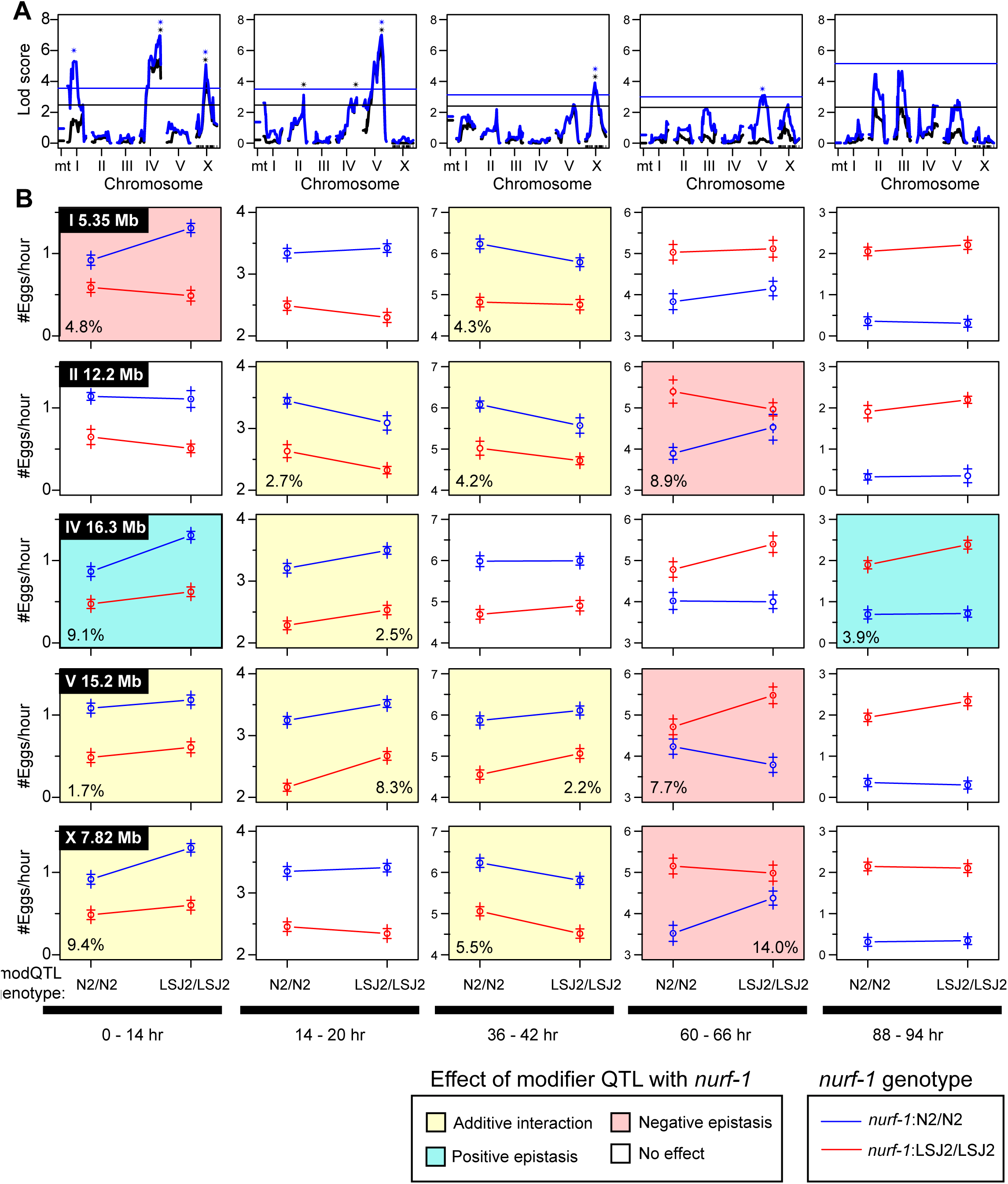
Additional modifier QTLs affect egg-laying in an age-dependent manner. **A**. QTL mapping using the *nurf-1* genotype as an additive (black) or interactive (blue) covariate. Black and blue stars indicate the five QTLs with genome-wide significance above 0.05. **B.** Average egg-laying rate of RILs partitioned and averaged by their genotype at *nurf-1* and one of the modifier QTLs at five time points. The modifier QTL genotype is indicated on the x-axis. The *nurf-1* genotype is indicated by the color of the lines. Non-white background coloring indicates a significant effect of the modifier QTL by ANOVA (p<0.05). Yellow background indicates a significant effect of the modifier QTL but no significant non-linear interaction with *nurf-1* by ANOVA (p < 0.05). Green background indicates a significant effect of the modifier QTL and a significant positive non-linear interaction with *nurf-1* by ANOVA (p < 0.05). Red background indicates a significant effect of the modifier QTL and a significant negative non-linear interaction with *nurf-1* by ANOVA (p < 0.05).

In order to analyze how these additional five modifier QTLs regulated the egg-laying rate at the five mapping time points, we segregated the 94 RIL lines based upon their *nurf-1* and modifier QTL genotype (**Figure 5B**). By segregating both genotypes, we could visually determine if any non-linear interactions (i.e. epistasis) existed between *nurf-1* and the five modifier loci. Additive interactions would result in two lines with identical slopes while non-linear interactions result in lines with different slopes. Visual inspection of these effect-size graphs indicated that both additive and non-linear effects were present. In order to formalize this analysis, we used ANOVA to determine (i) if the modifier QTL had a significant effect at the particular timepoint, (ii) whether there was a significant interaction term between *nurf-1* and the modifier variant, and (iii) the total amount of variance that the modifier QTL explained at that particular timepoint (**Figure 5B**). We considered two types of epistasis, positive and negative. Positive epistasis is defined as when the modifier variant has the same direction of effect in both *nurf-1* genotypes (e.g. the slope is positive for both lines but different magnitude). Negative epistasis is defined when the modifier variant has a different direction of effect in the different *nurf-1* genotypes (e.g. one slope is positive and the other slope is negative). This analysis indicated that the effect of these modifier QTLs were also complex. Each QTL was age dependent and showed non-linear interactions with the *nurf-1* locus. This mapping suggests that the egg-laying differences between N2 and LSJ2 are multigenic, involve extensive epistatic interactions, and are highly age-dependent.

### Modeling also predicts age-dependent epistasis at latter timepoints

Further inspection of the age-dependence and epistasis of the modifier QTLs with *nurf-1* revealed a number of interesting trends (**Figure 5B**). Epistasis was most likely to occur at the first and last two time points (7 out of 8 significant effects) and less likely to be observed at the second and third time point (1 out of 7 significant effects). While statistically significant, it is difficult to interpret epistasis and the first and last time point, as the rate of egg-laying converges on zero in some of these genetic backgrounds. We were more intrigued by the negative epistasis observed in three of the modifier QTLs at the 60 - 66 hour time point. This is the time point where we observed effect-size switching for *nurf-1* and we wondered whether the two features could be related. We decided to use additional modeling to determine if epistasis could arise through unequal use of sperm. We modeled a modifier QTL by assuming it changed the oocyte generation rate (k_o_) in an additive fashion (**Figure S6A**) and calculated the egg-laying rate predicted by this model for the four possible genotype combinations *(nurf-1* and the modifier QTL) (**Figure S6B**). These values were plotted at three time points in a similar manner as **Figure 5B** (**Figure S6C**).

This analysis demonstrated the potential role for a modifier QTLs to create negative epistasis, specifically at a time when we observe negative epistasis in our QTL mapping data. How does negative epistasis arise? The reason for this can be deduced from our previous observation that the effect size of the modifier QTL will switch signs due to unequal use of sperm in the two genetic backgrounds. However, the exact time this occurs is also dependent on the genetic background of the *nurf-1* locus. The period of time in between these two intersections is when negative epistasis is observed as the modifier has a positive effect in one *nurf-1* background but has switched to the negative effect in the other *nurf-1* background. After the sign-switching occurs for both genotypes, the modifier allele again has the same direction of effect in both *nurf-1* backgrounds. We refer to this time window as the sign epistasis zone (**Figure S6B**).

### Segregating the effects of the modifier QTLs onto k_o_ and S_0_

Finally, we decided to test whether we could use the macroscale modelling of the egg-laying process to improve our QTL mapping. Instead of performing QTL mapping on each of the five time points, we used this data to estimate a k_o_ and S_0_ for each RIL strain (assuming as before that k_c_ = 0 and k_f_ was the same for all RIL strains). The distribution of these fitted parameters are plotted in **Figure 6A and 6B**. We next performed QTL mapping to identify genetic regions that influence these rates. As expected, we identified a number of loci that we found from previous analysis. The QTL surrounding *nurf-1* was identified as a regulator of both the oocyte generation rate (k_o_) and the number of self-sperm (S_0_). The modifier QTLs on II and X regulated the oocyte generation rate but not the number of sperm number, while the QTL on V had an effect on sperm number but not the oocyte generation rate. Interestingly, this analysis also identified a new QTL on chromosome III as a regulator of sperm number (**Figure 6B**), suggesting that modeling of the egg-laying process not only helps us understand the modifier QTLs effect but also can help identify new QTL loci.

**Figure 6.**
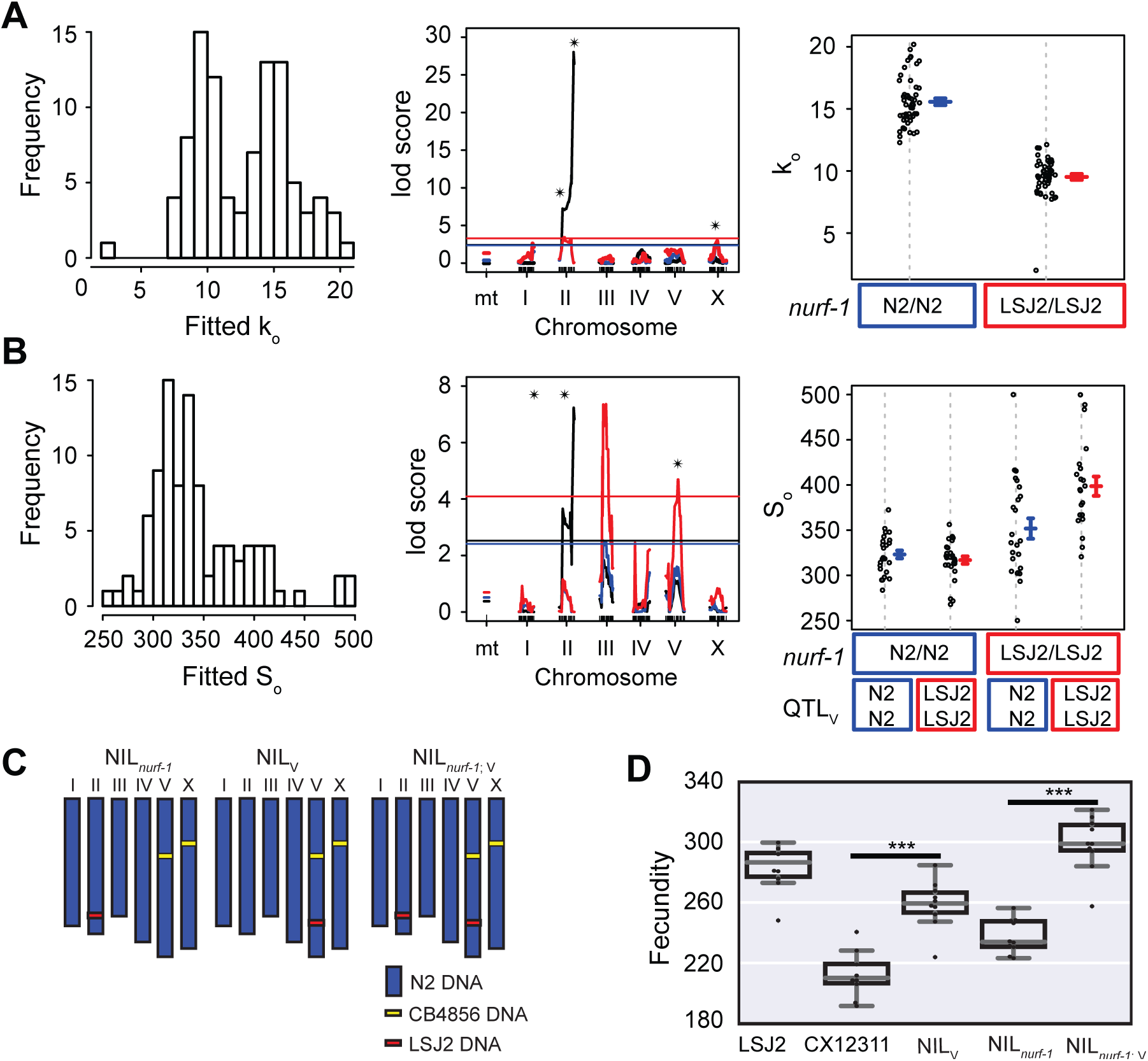
QTL mapping of parameters estimated for each RIL strain. **A and B.** A least-squares method was used to fit individual k_o_ and S_0_ values for all 94 RIL strains. **Left panel:** Histrogram of all k_o_ and S_0_ values from 94 RIL strains. **Middle panel:** QTL mapping of the k_o_ and S_0_ parameters. Black line indicates a one dimensional scan. We also used the *nurf-1* genotype as an additive (blue) or interactive (red) covariate. Black stars indicate QTLs with genome-wide significance above 0.05. **Right panel:** RIL strains segregated by their genotypes at the *nurf-1* locus (panel **A**) or their genotypes at the *nurf-1* and QTL_V_ loci (panel **B**). **C.** Schematic of NIL strains used for panel **D**. **D.** Total fecundity (# of eggs laid over the animal’s lifespan) of NIL strains and parental strains indicated on the x-axis.

In order to test these results, we utilized a NIL strain surrounding the QTL on V. We crossed the NIL_V_ strain to the NIL_*nurf-1*_ strain to create the double NIL_*nurf-1;*V_ strain (**Figure 6D**). This combination of strains allowed us to test the effect of the modifier QTL in both *nurf-1* genotypes. For each of these strains, we measured the total amount of progeny produced. These NIL strains showed that the QTL_V_ increased the brood size in both *nurf-1* backgrounds, but had a larger effect size in the LSJ2 *nurf-1* genotype (60 vs. 30) indicating epistasis does exist between these two loci. This qualitatively agrees with the QTL mapping data (**Figure 6B – right panel**) but discrepancies were observed between the magnitude of the total broodsizes, suggesting additional improvements to the model might be beneficial.

## DISCUSSION

Our results identified four new genetic loci that regulate reproductive rate and/or fecundity that arose and fixed following separation of the N2 and LSJ2 lineages. Polygenic traits are common in natural populations, but it was still surprising how rapidly a polygenic trait could evolve in laboratory conditions. Based upon minimum generation time, we estimate a maximum of 3900 generations separate the LSJ2 and N2 strains. We previously identified one of the causative genetic variants as a 60 bp deletion in *nurf-1*, which encodes a component of the NURF chromatin remodeling factor (29). This deletion arose and fixed in the LSJ2 lineage. Using competition experiments, we demonstrated this genetic variant was advantageous in the LSJ2 growth conditions, suggesting it was fixed by selection. The additional modifier variants could also be advantageous, which could explain their rapid fixation. However, due to the populations small effective population size and limited outcrossing, genetic draft and genetic drift are also relevant evolutionary forces for these strains. Additional work identifying the exact causative genetic variant will be necessary to determine the lineage it arose within and its fitness in the relevant conditions to determine if its fixation was caused by selection vs. genetic drift or draft. Unfortunately, these experiments are not trivial. Despite the inherent advantages to the system as compared to wild strains, each QTL contains a handful to dozens of potential causative genetic variants. The small effect-size of these variants also requires a large number of animals to be tested to obtain the statistical power to distinguish between strains with and without the variant. High-throughput and automated analysis of reproductive output would greatly aid this work.

The study of natural traits from a number of phyla has found a number of examples of biological epistasis (7, 12, 14, 39). Age can also act as an important covariate for genetic variation. The presence of epistasis and age-dependence can obscure the mapping between genotype and phenotype as the effect of causative genetic variants can wash out if mapping populations are not segregated appropriately by age or genetic background. Our results provide a framework to understand how age-dependence can also arise through emergent properties of a cellular network, which we believe to be the major scientific contribution of this work. Our work indicates that the complex seesaw of effects that *nurf-1* has on reproductive output can be explained using two major considerations: (1) a hormonally-mediated negative feedback loop linking sperm with oocyte maturation, and (2) the effect *nurf-1* has on the rate of oogenesis. Consequently, the molecular details of how *nurf-1* modifies protein and cellular function are unnecessary to explain its age-dependence. Our modeling experiments also demonstrated how sign epistasis could arise in an age-dependent manner strictly through age-independent changes in the oocyte production rate. The origin of genetic epistasis is often thought of in terms of biochemical properties of proteins, through physical interactions between two proteins, or through multiple changes to a parallel or linear signaling pathway (40-42). However, these mechanisms were typically identified through analysis of mutations or genetic perturbations generated in the laboratory that have a strong negative effect on fitness. It is interesting to us that such mechanisms don’t seem to be at play here. This is the second example we have identified in *C. elegans* of what we refer to as cellular epistasis (43) – i.e. the non-linear interactions are an emergent property of the functions of cellular networks as opposed to properties of molecular or biophysical interactions between proteins. It will be interesting to see how often cellular epistasis is responsible for genetic epistasis in natural traits.

While the *C. elegans* reproductive system is a special case, any negative feedback loops acting on a measurable trait could cause similar age-dependent changes. For example, negative autoregulatory feedback loops are common in transcription factors networks. Any genetic variants (either *cis* or *trans)* that increase the rate of transcription of one of these transcription factors will initially appear to have a positive effect-size on the mRNA levels of that gene. However, as the amount of protein product increases, the transcription factor will turn off the expression of its own mRNA. Since the amount of protein product will be higher in the strain with the higher initial expression, the transcription will turn off sooner in that strain, and the genetic variant will soon appear to have a negative effect on expression.

Our work provides an example of how multidisciplinary studies can be used tackle the genetic basis of complex traits – we leveraged quantitative genetics, detailed knowledge of the egg-laying process in *C. elegans,* and an existing macroscale model of egg-laying to make our conclusions.

## Acknowledgements

We thank the *Caenorhabditis* Genetics Center, Christian Frokjaer-Jensen, and Erik Jorgensen for strains; Erik Andersen, Joshua Weitz, and Greg Gibson for discussions; and Cori Bargmann, Greg Gibson, Todd Streelman, Hang Lu, and Andres Bendesky for helpful comments on the manuscript. This work was supported by NIH grants R21AG050304, R01GM114170, and the Ellison Medical Foundation (to P.T.M.).

## Author Contributions

E. E. L. and P.T.M. designed and interpreted experiments and wrote the paper. E. E. L. performed all egg-laying experiments and generated all NILs. E.E.L., R.P. and P.T.M. performed all QTL mapping. R.P and P.T.M. performed all modeling experiments. K.L.W performed all fertilized egg counts, large oocyte counts and sperm counts. K.L.W. and R.F.C. performed all mitotic cell counts. W.X. created *nurf-1* CRISPR-Cas9 strains. R.A.B. provided synthetic ascarosides.

## METHODS

### Strains

Strains were cultivated on agar plates seeded with *E. coli* strain OP50 at 20°C (44).

Strains used in this study are: N2, LSJ2, CX12311 *kyIR1(V, CB4856>N2); qgIR1(X, CB4856>N2).*

NIL strains used for this study are: PTM66 (NIL_*nurf-1*_) *kyIR87(II, LSJ2>N2); kyIR1(V, CB4856>N2); qgIR1(X, CB4856>N2),* PTM75 (NIL_V_) *kahIR2 (V,LSJ2>N2), kyIR1 (V, CB4856>N2); qgIR1(X, CB4856>N2),* PTM84 (NIL_*nurf-1*_, V) *kahIR6(IV,LSJ2>N2); kyIR87(II, LSJ2>N2); kyIR1(V, CB4856>N2); qgIR1(X, CB4856>N2)*

RIL strains used in this study are sequentially: CX12312 - CX12327, CX12346 - CX12377, CX12381 - CX12388, CX12414 - CX12437, and CX12495-CX12510

### Egg-laying assays

Egg-laying assays were performed as previously described (24).

For mating experiments in **figure 2E**, twelve N2 or CX12311 males were placed on experimental plates with six L4 hermaphrodites of interest for the first time point and mated hermaphrodites were then transferred to experimental plates without males. Successful mating events were validated via the observation of males in the F1 generation of mating plates.

### Immunofluorescence and Microscopy

Gonad immunofluorescence was performed on adult animals from each strain 24 and 48 hours after the L4 larval stage. Gonads were dissected in M9 containing 1% Tween and 1 mM levamisole and fixed in 2% paraformaldehyde via freeze cracking. Fixed gonads were stained with a 1:50 conjugated H3KS10p Alexa Fluor 488 antibody (Millipore Cat. 06-570-AF488) specific to mitotically dividing cells and 1.5 mg/mL of DAPI in Vectashield mounting medium (Vector Laboratories Cat. H-1200) (32, 33). Mitotic germline cells and germline stem cells were imaged and scored between the transition zone and the distal tip on an Olympus IX73 inverted microscope with 100×/1.40 UPlanSApo objective (Olympus) and a Hamamatsu Orca-flash4.0 digital camera. The distal tip cell (**Figure 2A**) is located at the tip of the gonad and releases a factor that promotes mitosis. The transition zone represents the location in the gonad where cells are beginning to undergo meiosis.

Embryo/oocytes/sperm/germline stem cells were quantified by fixing whole animals in 95% ethanol and staining nuclei with 1.5 mg/mL DAPI contained in Vectashield mounting medium. Embryos were scored using DIC on a 10x/0.30 UPlanFL N objective (Olympus). DAPI stained oocytes were imaged and scored between the spermatheca and posterior gonadal arm with a 40×/1.3 PlanApo objective (Olympus). Sperm and germline stem cells were identified using z-stack to capture multiple planes within each spermatheca or progenitor zone respectively. The transition zone was identified using morphological criteria based upon the crescent shape of the nucleus. All images and scoring were processed in ImageJ.

### Statistics

Significant differences between means were determined using a Mann-Whitney U test, a nonparametric test of the null hypothesis. For simplicity, a Bonferroni correction was used to modify the type I error rate to account for multiple testing. For each figure, we have listed the uncorrected p-value and the number of comparisons used for the Bonferroni correction in Supplemental Table 1. Both the non-parametric test and the Bonferroni correction should be conservative approximations of the true p-value. For the analysis of epistasis in Figure 5, we used ANOVA to calculate a F value and associated p-value using the fitqtl function in R/qtl. We calculated a single position for each of the five modifier QTLs to use for all five time points. All significant QTLs were simultaneously fit together for each of the five timepoints considering both their linear effect and their interaction with the *nurf-1* genoypte (i.e. y = QI +QII + Qnurf + QIV + QV + QX + QI*Qnurf + QII*Qnurf + QIV*Qnurf + QV*Qnurf + QX*Qnurf). The F value was calculated by dropping a single QTL at a time. Since most of the parameters were significant before multiple comparison testing, we used the Benjamini-Hochberg procedure to control for the false discovery rate.

### QTL Mapping

R/qtl was used to perform a one-dimensional scan using marker regression on the 192 markers (38). The significance threshold (p = 0.05) was determined using 1000 permutation tests. To identify modifier QTLs, the *nurf-1* marker was used as an additive and interactive covariate for additional one-dimensional scans, assuming a normal model. The significance threshold (p =0.05) for these two tests was determined using 1000 permutation tests.

### Egg-laying model

We adapted a previously published model (37), which simulates the egg-laying rate using the following set of equations:

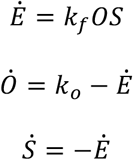

where *E* is fertilized eggs, *O* is oocytes, S is sperm, k_f_ is the rate of oocyte fertilization, k_o_ is the rate of ovulation, and k_c_ is the carrying capacity of the uterus. A dot indicates a time-derivative. We solved these ODEs numerically using a Dormand-Prince explicit solver or using an analytical solution (below). To calculate best fits, we assumed that the LSJ2, CX12311, RILs and NIL_*nurf-1*_ strains had unique values of k_o_, but shared the k_f_ parameters. These parameters were estimated using a Levenberg-Marquardt non-linear least squares algorithm. To calculate effect-size, we subtracted the two fits.

We used Mathematica to analytically solve the ODE equations:

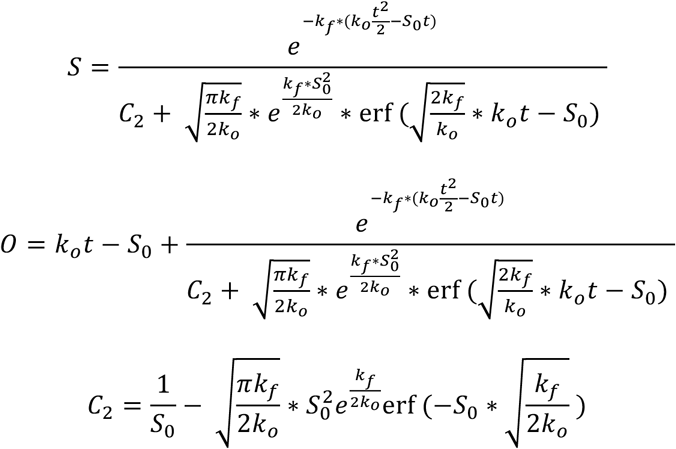

We used these equations to estimate the parameters using a Levenberg-Marquardt non-linear least squares algorithm.

### NIL strains

NILs were generated from previously existing RILs by backcrossing the chromosomal region of interest into a balancer strain containing a CX12311 background along with a fluorescent miniMos insertion near the QTL of interest. For ten generations, males that were heterozygous for the fluorescent marker (as determined by fluorescence intensity) were crossed to hermaphrodites of the balancer strain. On the 11^th^ generation, animals without the fluorescent marker were isolated. Genotyping of one to three markers within the candidate regions was used to confirm the successful introgression of LSJ2 DNA into CX12311. For each NIL strain, the starting RIL strain and the minMos balancer are listed in the table below:

| NIL strain | RIL strain | miniMos Balancer |
| --- | --- | --- |
| PTM66 | CX13248 | None; followed LSJ2 allele using PvuII-based genotyping |
| PTM75 | CX12361 | oxTi710 |

## REFERENCES

1. Wei WH, Hemani G, Haley CS. Detecting epistasis in human complex traits. Nat Rev Genet. 2014;15(11):722–33.

2. Phillips PC. The language of gene interaction. Genetics. 1998;149(3):1167–71.

3. Moore JH, Williams SM. Traversing the conceptual divide between biological and statistical epistasis: systems biology and a more modern synthesis. Bioessays. 2005;27(6):637–46.

4. Phillips PC. Epistasis--the essential role of gene interactions in the structure and evolution of genetic systems. Nat Rev Genet. 2008;9(11):855–67.

5. Brem RB, Storey JD, Whittle J, Kruglyak L. Genetic interactions between polymorphisms that affect gene expression in yeast. Nature. 2005;436(7051):701–3.

6. Deutschbauer AM, Davis RW. Quantitative trait loci mapped to single-nucleotide resolution in yeast. Nature genetics. 2005;37(12):1333–40.

7. Gerke J, Lorenz K, Cohen B. Genetic interactions between transcription factors cause natural variation in yeast. Science. 2009;323(5913):498–501.

8. Doebley J, Stec A, Gustus C. teosinte branched1 and the origin of maize: evidence for epistasis and the evolution of dominance. Genetics. 1995;141(1):333–46.

9. Kroymann J, Mitchell-Olds T. Epistasis and balanced polymorphism influencing complex trait variation. Nature. 2005;435(7038):95–8.

10. Rowe HC, Hansen BG, Halkier BA, Kliebenstein DJ. Biochemical networks and epistasis shape the Arabidopsis thaliana metabolome. The Plant cell. 2008;20(5):1199–216.

11. Wentzell AM, Rowe HC, Hansen BG, Ticconi C, Halkier BA, Kliebenstein DJ. Linking metabolic QTLs with network and cis-eQTLs controlling biosynthetic pathways. PLoS genetics. 2007;3(9):1687–701.

12. Huang W, Richards S, Carbone MA, Zhu D, Anholt RR, Ayroles JF, et al. Epistasis dominates the genetic architecture of Drosophila quantitative traits. Proceedings of the National Academy of Sciences of the United States of America. 2012;109(39):15553–9.

13. Mackay TF, Stone EA, Ayroles JF. The genetics of quantitative traits: challenges and prospects. Nature reviews Genetics. 2009;10(8):565–77.

14. Gaertner BE, Parmenter MD, Rockman MV, Kruglyak L, Phillips PC. More than the sum of its parts: a complex epistatic network underlies natural variation in thermal preference behavior in Caenorhabditis elegans. Genetics. 2012;192(4):1533–42.

15. Glater EE, Rockman MV, Bargmann CI. Multigenic natural variation underlies Caenorhabditis elegans olfactory preference for the bacterial pathogen Serratia marcescens. G3. 2014;4(2):265–76.

16. Carlborg O, Jacobsson L, Ahgren P, Siegel P, Andersson L. Epistasis and the release of genetic variation during long-term selection. Nature genetics. 2006;38(4):418–20.

17. Pettersson M, Besnier F, Siegel PB, Carlborg O. Replication and explorations of high-order epistasis using a large advanced intercross line pedigree. PLoS genetics. 2011;7(7):e1002180.

18. Cheverud JM, Vaughn TT, Pletscher LS, Peripato AC, Adams ES, Erikson CF, et al. Genetic architecture of adiposity in the cross of LG/J and SM/J inbred mice. Mammalian genome: official journal of the International Mammalian Genome Society. 2001;12(1):3–12.

19. Jarvis JP, Cheverud JM. Mapping the epistatic network underlying murine reproductive fatpad variation. Genetics. 2011;187(2):597–610.

20. Leamy LJ, Gordon RR, Pomp D. Sex-, diet-, and cancer-dependent epistatic effects on complex traits in mice. Frontiers in genetics. 2011;2:71.

21. Peripato AC, De Brito RA, Matioli SR, Pletscher LS, Vaughn TT, Cheverud JM. Epistasis affecting litter size in mice. Journal of evolutionary biology. 2004;17(3):593–602.

22. Francesconi M, Lehner B. The effects of genetic variation on gene expression dynamics during development. Nature. 2014;505(7482):208–11.

23. Cordell HJ. Detecting gene-gene interactions that underlie human diseases. Nature reviews Genetics. 2009;10(6):392–404.

24. Mackay TF. Epistasis and quantitative traits: using model organisms to study gene-gene interactions. Nature reviews Genetics. 2014;15(1):22–33.

25. McGrath PT, Rockman MV, Zimmer M, Jang H, Macosko EZ, Kruglyak L, et al. Quantitative mapping of a digenic behavioral trait implicates globin variation in C. elegans sensory behaviors. Neuron. 2009;61(5):692–9.

26. McGrath PT, Xu Y, Ailion M, Garrison JL, Butcher RA, Bargmann CI. Parallel evolution of domesticated Caenorhabditis species targets pheromone receptor genes. Nature. 2011;477(7364):321–5.

27. de Bono M, Bargmann CI. Natural variation in a neuropeptide Y receptor homolog modifies social behavior and food response in C. elegans. Cell. 1998;94(5):679–89.

28. Persson A, Gross E, Laurent P, Busch KE, Bretes H, de Bono M. Natural variation in a neural globin tunes oxygen sensing in wild Caenorhabditis elegans. Nature. 2009;458(7241):1030–3.

29. Large EE, Xu W, Zhao Y, Brady SC, Long L, Butcher RA, et al. Selection on a Subunit of the NURF Chromatin Remodeler Modifies Life History Traits in a Domesticated Strain of Caenorhabditis elegans. PLoS Genet. 2016;12(7):e1006219.

30. Duveau F, Felix MA. Role of pleiotropy in the evolution of a cryptic developmental variation in Caenorhabditis elegans. PLoS biology. 2012;10(1):e1001230.

31. Dalfo D, Michaelson D, Hubbard EJ. Sensory regulation of the C. elegans germline through TGF-beta-dependent signaling in the niche. Curr Biol. 2012;22(8):712–9.

32. Hendzel MJ, Wei Y, Mancini MA, Van Hooser A, Ranalli T, Brinkley BR, et al. Mitosis-specific phosphorylation of histone H3 initiates primarily within pericentromeric heterochromatin during G2 and spreads in an ordered fashion coincident with mitotic chromosome condensation. Chromosoma. 1997;106(6):348–60.

33. Kalchhauser I, Farley BM, Pauli S, Ryder SP, Ciosk R. FBF represses the Cip/Kip cell-cycle inhibitor CKI-2 to promote self-renewal of germline stem cells in C. elegans. EMBO J. 2011;30(18):3823–9.

34. Schafer WF. Genetics of egg-laying in worms. Annual review of genetics. 2006;40:487–509.

35. Miller MA, Nguyen VQ, Lee MH, Kosinski M, Schedl T, Caprioli RM, et al. A sperm cytoskeletal protein that signals oocyte meiotic maturation and ovulation. Science. 2001;291(5511):2144–7.

36. Miller MA, Ruest PJ, Kosinski M, Hanks SK, Greenstein D. An Eph receptor sperm-sensing control mechanism for oocyte meiotic maturation in Caenorhabditis elegans. Genes & development. 2003;17(2):187–200.

37. McMullen PD, Aprison EZ, Winter PB, Amaral LA, Morimoto RI, Ruvinsky I. Macro-level modeling of the response of C. elegans reproduction to chronic heat stress. PLoS computational biology. 2012;8(1):e1002338.

38. Broman KW, Wu H, Sen S, Churchill GA. R/qtl: QTL mapping in experimental crosses. Bioinformatics. 2003;19(7):889–90.

39. Hemani G, Shakhbazov K, Westra HJ, Esko T, Henders AK, McRae AF, et al. Detection and replication of epistasis influencing transcription in humans. Nature. 2014;508(7495):249–53.

40. Gertz J, Gerke JP, Cohen BA. Epistasis in a quantitative trait captured by a molecular model of transcription factor interactions. Theor Popul Biol. 2010;77(1):1–5.

41. Harms MJ, Thornton JW. Evolutionary biochemistry: revealing the historical and physical causes of protein properties. Nature reviews Genetics. 2013;14(8):559–71.

42. Lehner B. Molecular mechanisms of epistasis within and between genes. Trends in genetics: TIG. 2011;27(8):323–31.

43. Greene JS, Dobosiewicz M, Butcher RA, McGrath PT, Bargmann CI. Regulatory changes in two chemoreceptor genes contribute to a Caenorhabditis elegans QTL for foraging behavior. Elife. 2016;5.

44. Brenner S. The genetics of Caenorhabditis elegans. Genetics. 1974;77(1):71–94.

